# The Impact of Pathogenic and Artificial Mutations on Claudin-5 Selectivity from Molecular Dynamics Simulations

**DOI:** 10.1101/2023.01.30.526089

**Authors:** Alessandro Berselli, Giulio Alberini, Fabio Benfenati, Luca Maragliano

**Author notes:** Correspondence: F.B. -, L.M. These authors contributed equally to this work (Alessandro Berselli, Giulio Alberini).

## Abstract

Tight junctions (TJs) are multi-protein complexes at the interface between adjacent endothelial or epithelial cells. In the blood-brain barrier (BBB), they are responsible for sealing the paracellular spaces and their backbone is formed by Claudin-5 (Cldn5) proteins. Despite the important role in preserving brain homeostasis, little is known on how Cldn5 oligomers assemble. Different structural models have been suggested, where Cldn5 protomers from opposite cells associate to generate paracellular pores that do not allow the passage of ions or small molecules. Recently, the first Cldn5 pathogenic mutation, G60R, was identified and shown to induce anion selectivity in the BBB TJs. This offers an excellent opportunity to further assess the structural models. In this work, we performed umbrella sampling molecular dynamics simulations to study the permeation of single Na^+^, Cl^−^ and H_2_O through two distinct G60R Cldn5 paracellular models. Only one of them, called Pore I, reproduces the functional modification observed in the experiments, displaying a free energy (FE) minimum for Cl^−^ and a barrier for Na^+^ at the central constriction, consistent with the formation of an anionic channel. To further test the validity of the model, we performed the same calculations for the Q57D and the Q63D mutants, which affect two side-chains in the constriction site. In particular, Q57 is conserved among various Cldns, with few exceptions such as the two cation permeable homologs Cldn15 and Cldn10b. In both cases, we obtain that the FE profiles are modified with respect to the wild-type system, facilitating the passage of cations. Our calculations are the first *in-silico* description of the effect of a Cldn5 pathogenic mutation, and provide a further assessment of the Pore I model for Cldn5-based TJ architectures, yielding new atom-detailed insight on the selective permeability of the paracellular spaces in BBB.

## Introduction

The blood-brain barrier (BBB) is a highly selective interface that separates the capillary blood flow from the brain parenchyma, and it is composed of brain endothelium, pericytes and astrocytes. Brain endothlial cells are tightly bound at their lateral membranes by multimeric protein complexes named tight junctions (TJs) (1–3), that are quite impermeable to solutes. TJ proteins are organized in continuous networks of transmembrane strands, and are responsible for strictly regulating the diffusion of solutes through the intercellular space between cells, named the paracellular space. Understanding their molecular mechanisms and dysfunctions has a very high clinical potential (4, 5).

The protein Claudin 5 (Cldn5) is the most abundant component of the BBB TJ strands and it serves as their backbone. To generate TJs, Cldn5 proteins assemble via *cis*-(intracellular) interactions within each endothelial cell membrane and via *trans-* (intercellular) interactions across adjacent cells. The Cldn5 topology is characterized by a four transmembrane helix bundle (TM1-4) with two extracellular loops (ECL1-2), cytoplasmic N- and C-terminal residues, and an intracellular loop (2). The physiological function of Cldn5 as a regulator of paracellular transport makes it a relevant target in the development of novel strategies to deliver drugs directly to the brain. However, structure-based approaches are still limited by our incomplete understanding of how Cldn5 generates high-order complexes.

Recently, different structural models were proposed as building blocks of Cldn5 paracellular architectures (6–8). Two of them (6, 7), usually called Pore I and Pore II models, give rise to a pore cavity in the paracellular space, and their validity was investigated also for other Cldns homologous to Cldn5 and expressed in non-neuronal tissues. (9–22) While Pore I was first proposed for Cldn15 based on experimental observations (10), Pore II was originally suggested by computational studies (7). Albeit different, both configurations are formed by two *cis-* dimers that associate paracellularly to generate a tetramer encompassing a pore. Remarkably, in the case of Cldn5, molecular dynamics (MD) simulations showed that both pores do not allow the permeation of ions or small molecules through the paracellular space, in line with the protein’s known barrier function (23, 24). In our previous work (24), we calculated the free energy (FE) of single ions/water molecules translocating across the two Cldn5 models, revealing that both structures oppose to ionic permeation, but are permissive to water.

The first *de novo* Cldn5 mutation linked to a neurological condition was recently reported (25). The variation, G60R, is located in the ECL1 domain and is associated with two unrelated cases of alternating hemiplegia with microcephaly. Experiments showed that, while the mutation does not alter the formation of TJs by Cldn5, it strongly affects its barrier function, by inducing high permeability for Cl^−^ and low permeability for Na^+^. The discovery and the characterization of this mutation provides an excellent opportunity to further assess the two aforementioned structural models of Cldn5 complexes. Here, we performed umbrella sampling (US) MD simulations to calculate the FE of permeation of single Na^+^, Cl^−^ and H_2_O molecules through the two models for Cldn5^G60R^, and compared results with those of the native, wild type, protein Cldn5^WT^ (24). Notably, only Pore I, where the arginine side-chain is located near the constriction site at the center of the pore, reproduces the experimentally observed change in the ion selectivity induced by the mutation. Indeed, the mutated Pore I Cl^−^ and Na^+^ FE profiles along the channel axis are characterized by an attractive minimum and a repulsive barrier, respectively, thus suggesting the formation of an anionic channel.

With these results at hand, and to further test the validity of the Pore I model, we performed the same calculations after the mutation of two glutamine residues, also located at the central constriction site of the Cldn5 structure, into aspartate (Q57D and G63D, respectively). Q57 is conserved among various Cldns, with some relevant exceptions including Cldn15 and Cldn10b, where the residue is replaced by an aspartate that plays a pivotal role in conferring ionic selectivity to the paracellular channels. As expected, we observed that both mutations modify the FE profile, making it attractive to cations and repulsive to anions.

This study provides an *in-silico*, atom-level rationale explaining the effect of a pathogenic mutation on the Cldn5mediated BBB paracellular properties. Altogether, our findings further validate the Pore I model for the paracellular arrangement of Cldn5, revealing it as the sole consistent with all the available experimental observations. By identifying the protein-protein surface at the core of TJ Cldn5 assemblies, our computational investigation provides a step forward in the establishment of a robust basis for rational design of small molecules capable of modulating the BBB’s paracellular permeability in a safe and reversible manner.

## Materials and Methods

### The configurations of the two Cldn5 models and MD parameters

In order to perform the simulations for the mutated Cldn5 systems, we used two equilibrated wild type (WT) configurations of the single pore models (Pore I and Pore II), as described in our previous work (24). In addition to the insertion of the mutations, we added counterions in both solvent exposed regions, in order to re-neutralize the system. Briefly, each model includes four Cldn5 protomers (two for each *cis-* configuration), two membrane bilayers formed by 1-palmitoyl-2-oleoyl-SN-glycero-3phosphocholine (POPC) molecules, water molecules and ions. The G60R mutation was included in each Cldn5 subunit with the help of UCSF Chimera (26), and the initial configurations of the R side-chains were selected from the Dunbrack rotamers library (27). The resulting systems were refined with GalaxyRefineComplex (28, 29), embedded in a double POPC bilayer, solvated with explicit water molecules and neutralized with counterions. CHARMM hydrogen atoms were added with CHARMM-GUI (30–32) and disulfide bonds were assigned to the couples of conserved cysteine residues, according to the information provided in the Cldn PDB ID: 4P79 structure.

As in our previous works (11, 24), we used the NAMD software (33). All the simulations were performed using an *NpT* ensemble (*p* = 1 atm, *T* = 310 K), employing the Nosé-Hoover Langevin piston method (34, 35) to maintain the pressure at 1 atm, and a Langevin thermostat at 310 K. Following the details of the CHARMM-GUI configuration files for NAMD, the oscillation period of the piston was set at 50 fs, and the damping time scale at 25 fs. The Langevin thermostat was set with a damping coefficient of 1 ps^−1^.

The CHARMM36m (36–38)/CHARMM36 39) parameters were used for the protein and lipids, respectively, together with the TIP3P model for water molecules (40) and the associated ionic parameters with NBFIX corrections (41–43). Electrostatic and van der Waals interactions were calculated with a cutoff of 12 Å as prescribed by the CHARMM force field. A smoothing decay was applied starting to take effect at 10 Å. Hexagonal periodic boundary conditions implemented in NAMD were adopted to limit the size of the system. Long range electrostatic interactions were calculated using the Particle Mesh Ewald (PME) algorithm (44), adopting a spline interpolation order 6. A maximum space between grid points of 1.0 Å was used, as implemented in CHARMM-GUI. All covalent bonds involving hydrogen atoms were constrained using the SHAKE/SETTLE algorithms (45, 46), in order to use a time-step of 2 fs. To ensure maximum accuracy, electrostatic and van der Waals interactions were computed at each simulation step (24).

Consistent with the protocol of Ref. 24, an initial minimization and an additional 30-ns-long equilibration with progressive release of positional restraints, were performed to achieve a stable starting configuration for US runs.

1D-US simulations with harmonic biases were performed with the Collective Variables (Colvar) module (47), using the projection of the ion position onto the axis of the paracellular channel as single collective variable (CV). The Weighted Histogram Analysis Method (WHAM) was used to reconstruct the 1D-FE, using the code from the Grossfield group, available at http://membrane.urmc.rochester.edu/content/wham.

During US simulations, we employed harmonic restraints for the C_*α*_ atoms of the same set of residues introduced in Ref. 24, in order to avoid orientational and translational displacements of each paracellular model, that could perturb the FE calculations.

### Umbrella sampling simulations

We performed 1D-FE calculations of ion/water permeation through the paracellular cavities, following the same protocol used in our previous works (12, 13, 24). In umbrella sampling (US) calculations (48), a harmonic potential energy term is added to the CHARMM terms of the MD potential to ensure efficient sampling along the chosen CV in different independent simulations (named *windows*). In the case of our 1D CV, indicated as *z*, these potentials are expressed as:

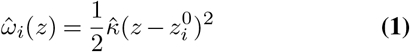

where 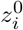 indicates the value at which the CV is restrained in window *i* and 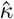 is a constant. For an appropriate choice of 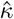, a sufficient overlap between the sampled distributions of adjacent windows is expected and the final, full FE calculations can be obtained by employing WHAM (49, 50).

We performed US simulations with the harmonic restraints of Eq. (1) and 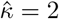. We employed a total of 60 windows with a uniform spacing of 1 Å. The time length required for convergence for each window is shown in **Table 1**. For each window, the frames from the first ns were removed and not used for analysis. To optimize the use of parallel computational resources, we prepared a number of starting conformations where the tested ion or the H_2_O oxygen atom was swapped with equilibrated water molecules, for the whole length of the paracellular axis. Ions not involved in the CV definition were excluded from the paracellular cavity by applying two half-harmonic potentials, one for each entrance, characterized by an elastic constant of 10 kcal/(mol · Å^2^.

**Table 1.** Summary of the MD simulations performed in this work.

| Model | System | Permeating ion | Time (ns) |
| --- | --- | --- | --- |
| Pore I | G60R | Na<sup>+</sup> | 10 × 60 = 600 |
| Pore I | G60R | Cl<sup>-</sup> | 40 × 60 = 2400 |
| Pore I | G60R | H<sub>2</sub>O | 10 × 60 = 600 |
| Pore II | G60R | Na<sup>+</sup> | 10 × 60 = 600 |
| Pore II | G60R | Cl<sup>-</sup> | 30 × 60 = 1800 |
| Pore II | G60R | H<sub>2</sub>O | 10 × 60 = 600 |
| Pore I | Q57D | Na<sup>+</sup> | 20 × 60 = 1200 |
| Pore I | Q57D | Cl<sup>-</sup> | 10 × 60 = 600 |
| Pore I | Q57D | H<sub>2</sub>O | 10 × 60 = 600 |
| Pore I | Q63D | Na<sup>+</sup> | 30 × 60 = 1800 |
| Pore I | Q63D | Cl<sup>-</sup> | 20 × 60 = 1200 |
| Pore I | Q63D | H<sub>2</sub>O | 10 × 60 = 600 |

In addition, the displacement of the ion orthogonal to the pore axis is confined within a disk of radius *r*_0_ + *δ*, where *r*_0_ is the pore radius as determined by the HOLE program (51, 52) and *δ* = 2 Å, by means of a force constant of 2 kcal/mol (24).

For WHAM analysis, we used 600 bins, and a tolerance of 0.0001. Furthermore, we used bootstrapping to evaluate the statistical uncertainty at each bin, using 100 bootstrap trials (12). Overall, the US calculations are based on a cumulative production of 12, 6 *μ*s.

### Structural analysis

All the MD trajectories were visualized and analyzed using UCSF Chimera (26) (www.cgl.ucsf.edu/chimera/), the NAMD-COLVAR module (47) and VMD (53) (www.ks.uiuc.edu/Research/vmd/) with Tcl scripts.

### Alignment of sequences

The alignment of multiple Cldn sequences was performed using the Clustal Omega program (54, 55) available in Jalview (56).

## Results

This section is organized into three paragraphs. In the first one, we show a sequence alignment among various Cldn sequences to visualize the positions of the three mutations investigated in this work. In the second, we report and discuss the FE profiles obtained from the two G60R pore configurations, extracted from the equilibrated structures of Pore I and Pore II. The resulting thermodynamic calculations are then compared with the results of the associated WT configurations published in Ref. 24. In the last part, we discuss the FE profiles of the Q57D and Q63D mutants of Pore I and compare the calculations with those of the WT model.

### The natural and artificial mutations involve highly conserved residues in the Cldn family

As in all other Cldns, the ECL1 and ECL2 of human Cldn5 (hCldn5) are organized in a *β*-sheet formed by five antiparallel strands, named *β*1 to *β*5. The first four strands belong to the ECL1 domain and the fifth one to the ECL2 domain. The ECL1 domain is stabilized by a conserved disulfide bond formed by two Cys residues belonging to *β*3 and *β*4. An additional extracellular helix (named ECH) is observed between *β*4 and the second transmembrane helix (TM2). In **Fig. 1**, we show a multisequence alignment of a selection of human Cldn proteins. Interestingly, the G60 residue is conserved among all these sequences, with the exception of Cldn8 and Cldn17, and it is located in a region where mutations were predicted to be deleterious (25).

**Fig. 1.**
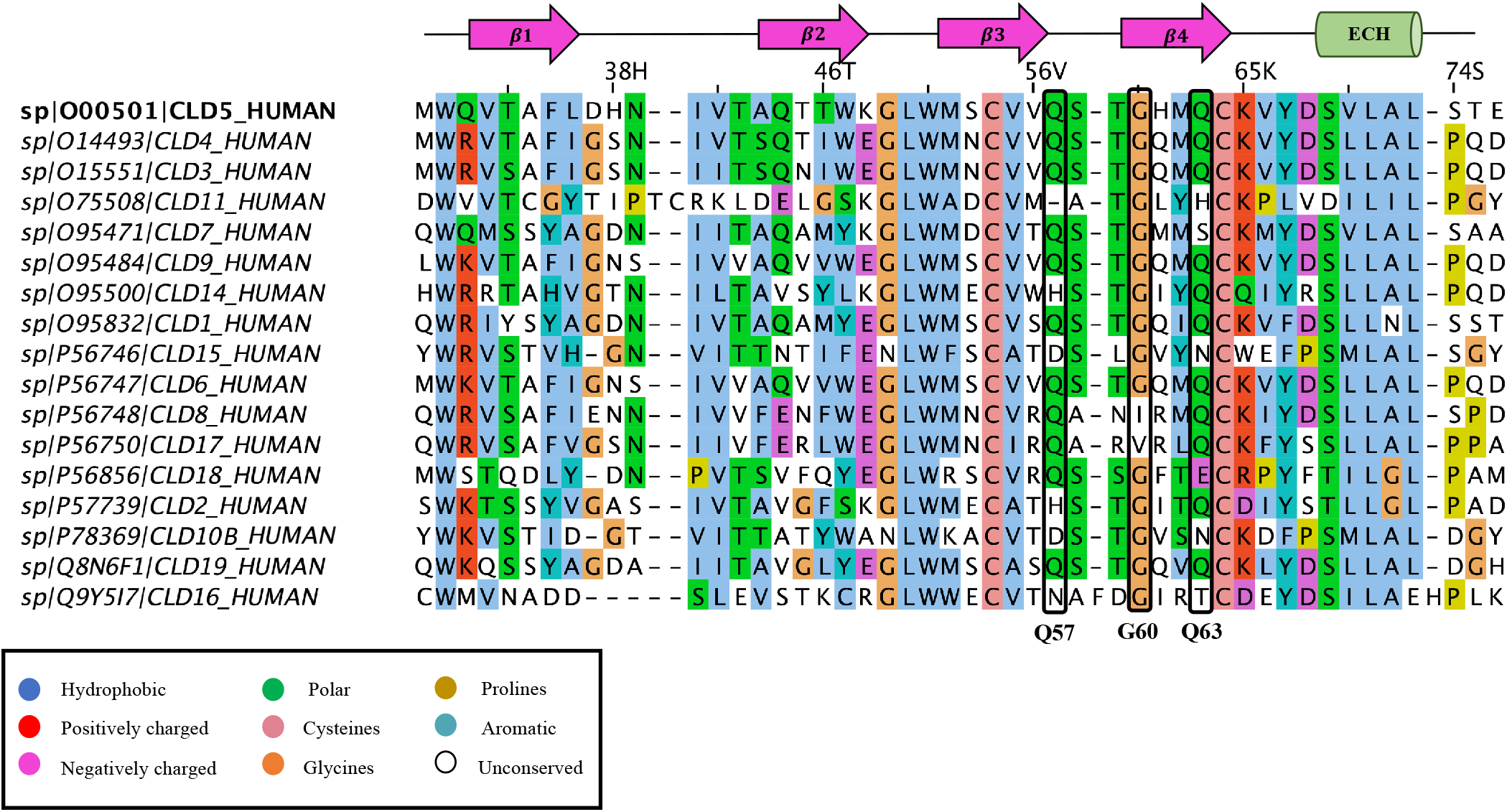
Multiple sequence alignment of human Cldn5 and other human Cldn proteins (ECL1 region only). This analysis includes classic Cldns (1–10, 14, 15, 17, 19) and two non-classic Cldns (11, 18). Each sequence is identified by the following entry: sp / UNIPROT (https://www.uniprot.org) ID / Cldn name. Colours are assigned according to the legend included in the picture. The numbering above the sequences is related to the Cldn5 protein. Secondary structure elements associated with the ECL1 topology are shown above the sequences as arrows (*β*1 to *β*4 strands) or cylinders (ECH *α*-helix). The Cldn5 residues corresponding to the mutations studied in this article (Q57, G60, Q63) are indicated by black rectangles.

The other two mutants (Q57D and Q63D) affect residues located in the same portion of the ECL1 domain, between *β*3 and *β*4 (Q57), and on *β*4 (Q63). In the cation-selective, channel-forming, Cldns 15 and Cldn10b proteins, an aspartate is found at the position equivalent to Cldn5 Q57(2), which prompted us to investigate the effect of the Q57D variant in the Cldn5 pore model.

### The G60R mutation induces an anion-selective channel in Pore I but not in Pore II

In order to perform the FE calculations of ion/water translocation through the mutated paracellular structures, we inserted the G60R variant in the WT configurations of both pore models. Due to the different orientation of the monomers, the mutated residues are located near the middle of the cavity in Pore I, and at the end-points of the cavity in Pore II (**Fig. 2**). Remarkably, however, in both structures the four mutated side-chains point outside the pore vestibule. Thus, in these single-pore Cldn5 models, it is expected that the G60R mutation influences ion translocation via electrostatic interactions only, excluding steric hindrance effects.

**Fig. 2.**
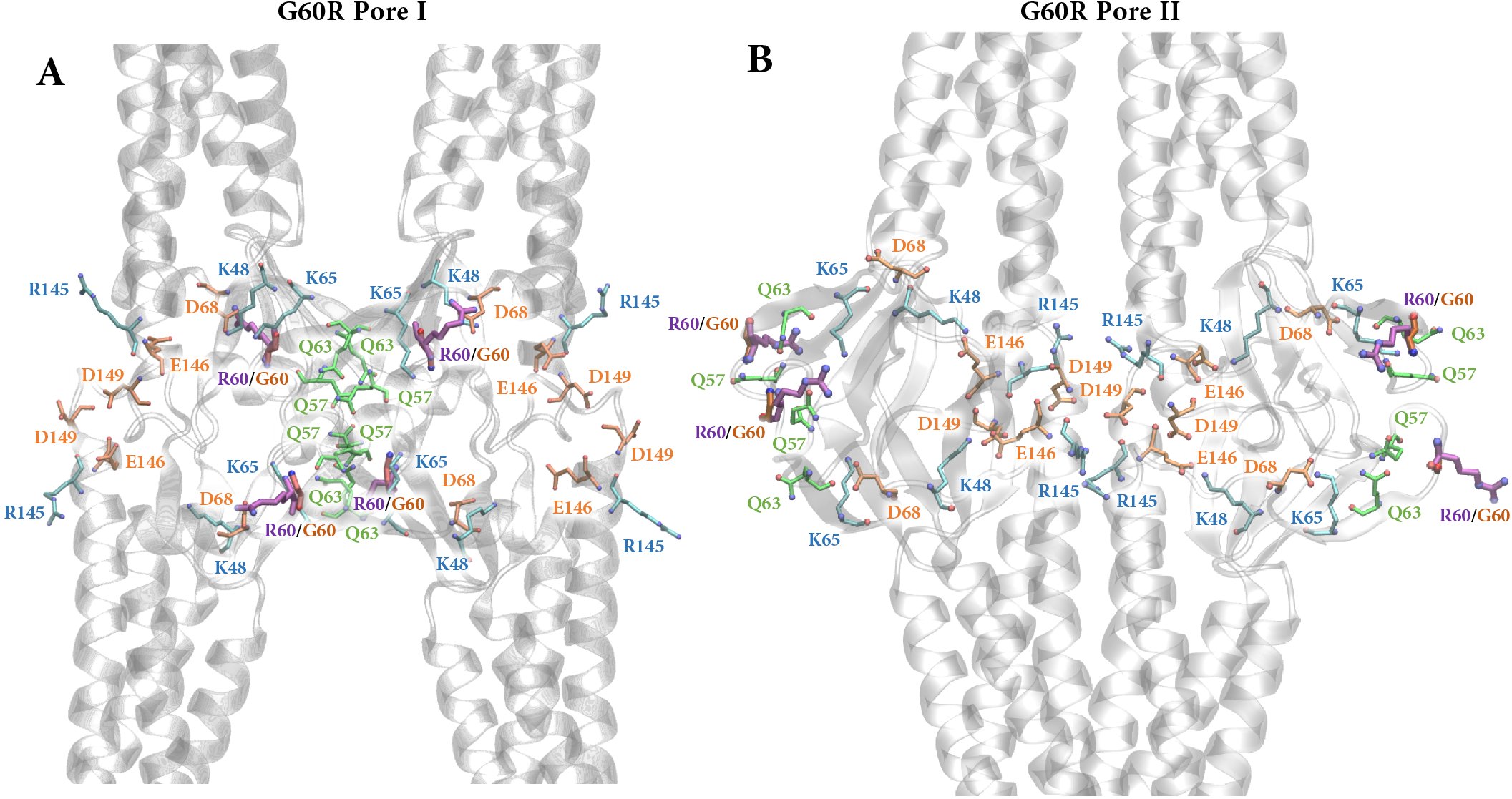
Equilibrated structures of G60R Cldn5 Pore I/II. The four protomers are represented as gray ribbons. The side-chains of the pore lining residues are represented as sticks. The mutated G60R side-chains are colored in purple.

The FE profiles for the G60R Cldn5 structures, calculated via US, are reported in **Fig. 3**, and compared with analogous profiles for the WT Cldn5 pores as calculated by us in Ref.24 (data reproduced with permission from ACS). Results show that the two models remain permeable to water after the insertion of the mutation, which induces only a small FE barrier in proximity of the mutated residues in Pore I (**Fig. 3A,B**).

**Fig. 3.**
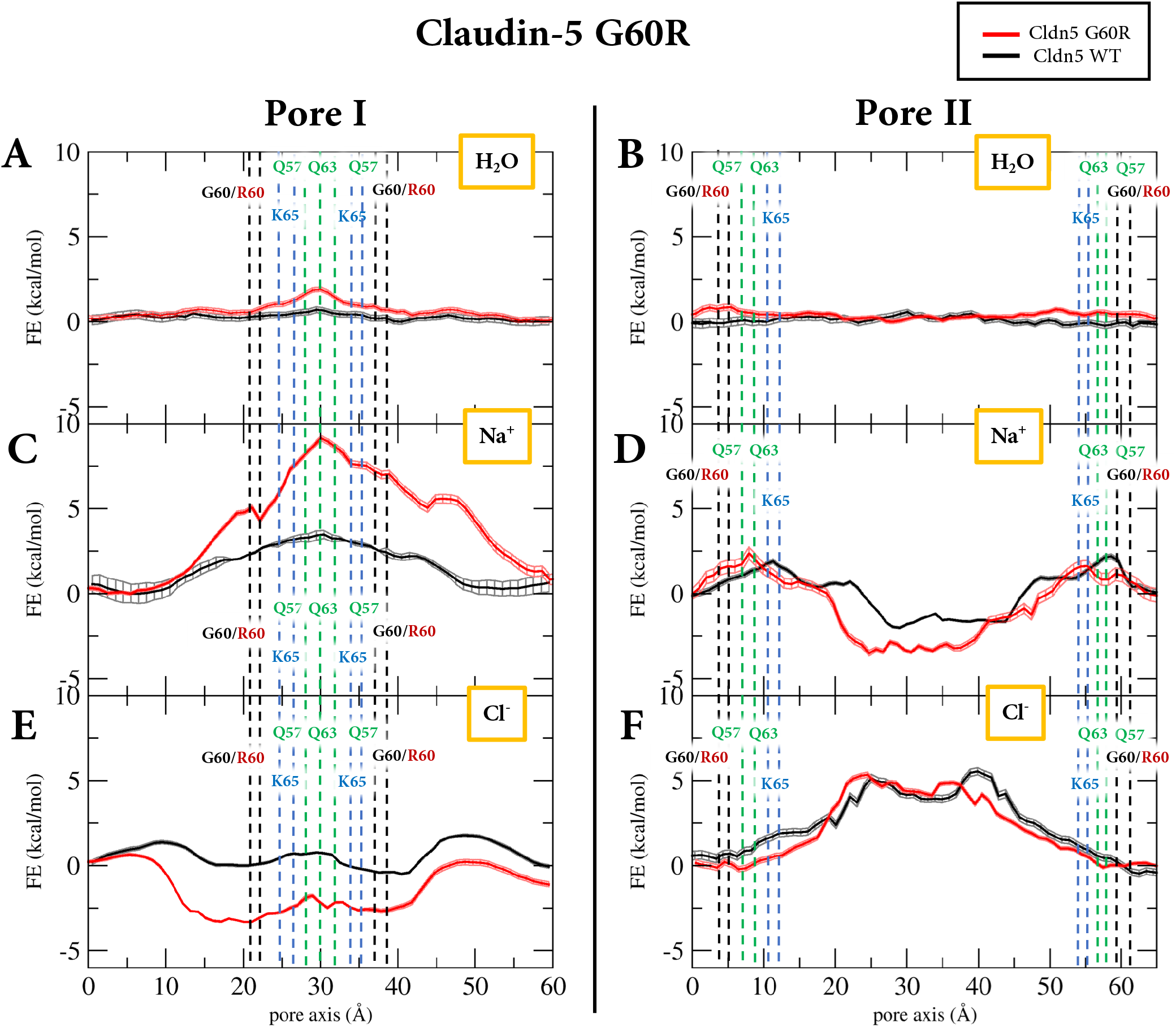
FE profiles for H_2_ O, Na^+^ and Cl^−^ translocation across the pore axis of both Pore I and Pore II systems. Red and black lines represent values for the G60R mutant and the WT, respectively (WT data reproduced with permission from Ref. 24, Copyright 2022 American Chemical Society). The positions of the C_*α*_ atoms of Q57, G60R, Q63 and K65 are reported as dashed lines.

Water translocation across the BBB is known to be characterized by limited exchange rate (57–59) and, to the best of our knowledge, there is no evidence to rule out water exchange through the Cldn5 paracellular pathways. Indeed, all computational investigations of the WT Cldn5 pores performed so far revealed water permeable cavities. (7, 23, 24).

On the other hand, Na^+^ translocation is affected in the Cldn5^G60R^ Pore I (**Fig. 3C**), where the four added arginine side-chains cause an increase of the central FE peak from 2.5-3 kcal/mol to 8.5-9 kcal/mol. In Pore II, the difference between the WT and the Cldn5^G60R^ Na^+^ FE profiles lies mainly at the center of the pore, where the mutations deepen the energy minimum by about 1-2 kcal/mol, while the barriers at the pore’s end-points are essentially preserved (**Fig. 3D**), in line with the location of the substituted arginine side-chains. In the case of Cl^−^, the mutated Pore I displays an FE minimum deeper than the WT of approximately −2.5 kcal/mol for 10 Å ≤ *z* ≤ 45 Å (**Fig. 3E**), consistent with the location of the arginines and with the function of an anionic channel. Conversely, the FE profile of Cl^−^ translocation in the G60R Pore II is essentially the same as in the WT configuration (i.e., a repulsive barrier; **Fig. 3F**), as a consequence of the different position of the mutated residues along the main axis of the paracellular cavity.

### The Q57D and Q63D mutations convert Pore I into a cationic paracellular channel

In order to further investigate the effect of pore-lining mutations in Pore I, we calculated the water/ion translocation FE profiles for the artificial mutants Q57D and Q63D. The locations of these residues are reported in **Fig. 4**. The glutamine in position 57 is highly conserved among Cldn proteins, with the notable exception of the two cation permeable Cldn15 (D55) and Cldn10b (D56) (2, 60). Results published in Ref.s 61, 62 extend the validity of the Pore I model to these members of the Cldn family, suggesting that they attract cations through the paracellular space thanks to the formation of a tetra-aspartate cage formed by the D55/D56 side-chains at the pore constriction. Q63 is also highly conserved in Cldns, but it is substituted by an asparagine in Cldn15 and in Cldn10b. Based on these observations, we asked ourselves wheter point mutations resulting in the insertion of negative charges in this critical region could transform the impermeable Cldn5^WT^ pore to an artificial paracellular cation channel. We thus generated the Cldn5^Q57D^ and Cldn5^Q63D^ mutants and studied the effect of the mutation on water and ion translocation.

**Fig. 4.**
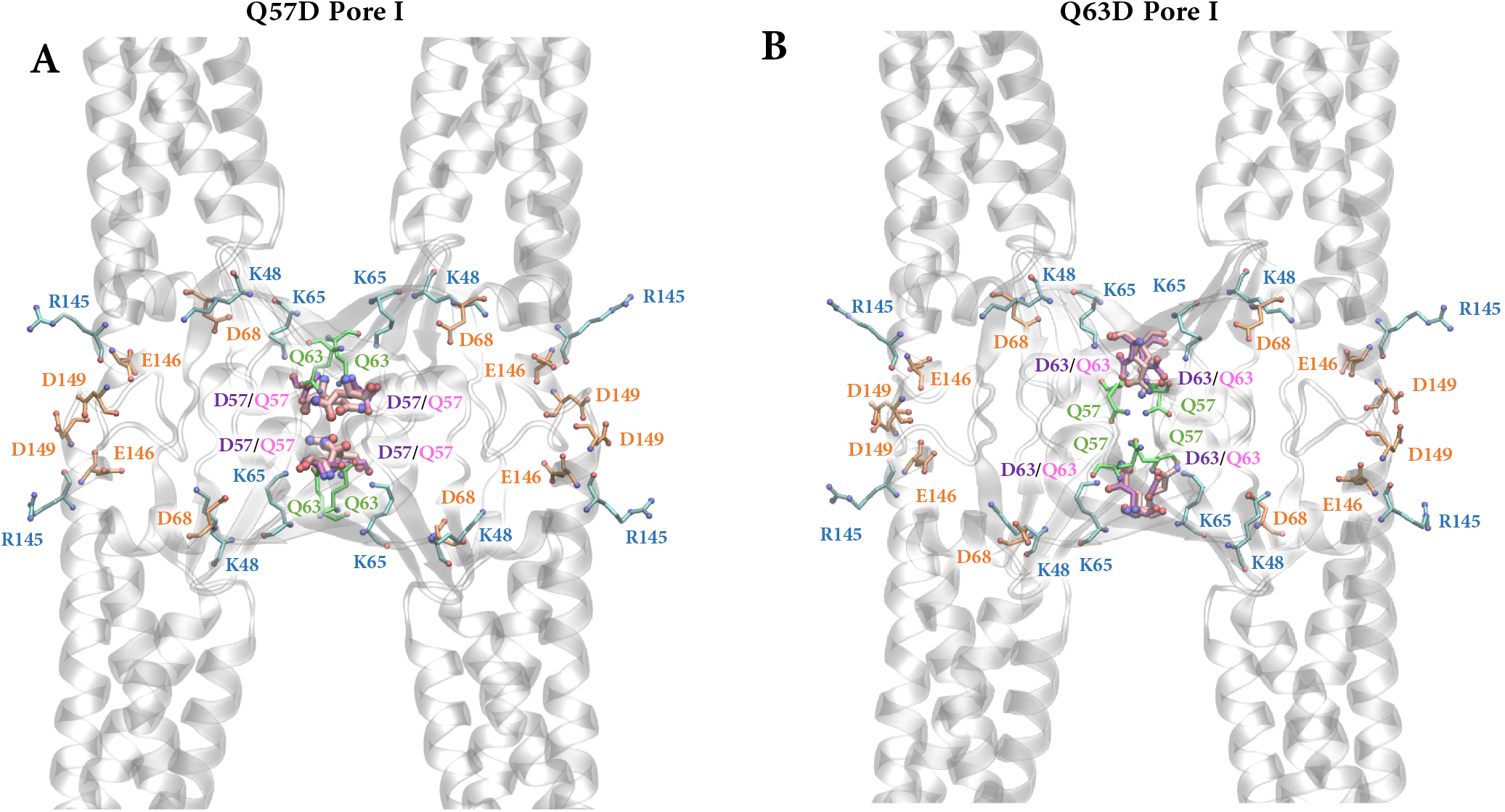
Representative structures of Pore I Q57D and Q63D configurations. The four Cldn5 proteins are represented as gray ribbons. The pore lining residues are reported, with their side-chains represented as sticks. The mutated Q57D and Q63D side-chains are colored in purple.

Our results based on US simulations of single Na^+^, Cl^−^ and H_2_O through the Cldn5^Q57D^ and Cldn5^Q63D^ Pore I configurations are shown in **Fig. 5**, and compared again with the WT system. Both mutations allow the structure to remain permissive to water paracellular diffusion **(Fig. 5A,B)**. As for Na^+^ translocation, they abolish the FE barrier in the central constriction seen in the WT system and cause the formation of a minimum. While the FE minimum of the Q63D mutation is localized in a narrow region of the paracellular axis (at 25 Å ≤ *z* ≤ 30 Å) and is modestly deep (~ 2 kcal/mol, **Fig. 5D)**, the Q57D substitution induces a broader (20 Å ≤ *z* ≤ 40 Å) and deeper minimum (~ 5 kcal/mol, **Fig. 5C)**, confirming the paramount role of this residue in the regulation of the paracellular traffic. Finally, both mutations also modify the FE Cl^−^ translocation profiles, causing the appearance of a central FE peak at ~ 7.5 kcal/mol **(Fig. 5E,F)**. Remarkably, the results for Na^+^ and Cl^−^ in the Q57D Pore I are consistent with our previous calculations for Cldn15 in the same configuration (12).

**Fig. 5.**
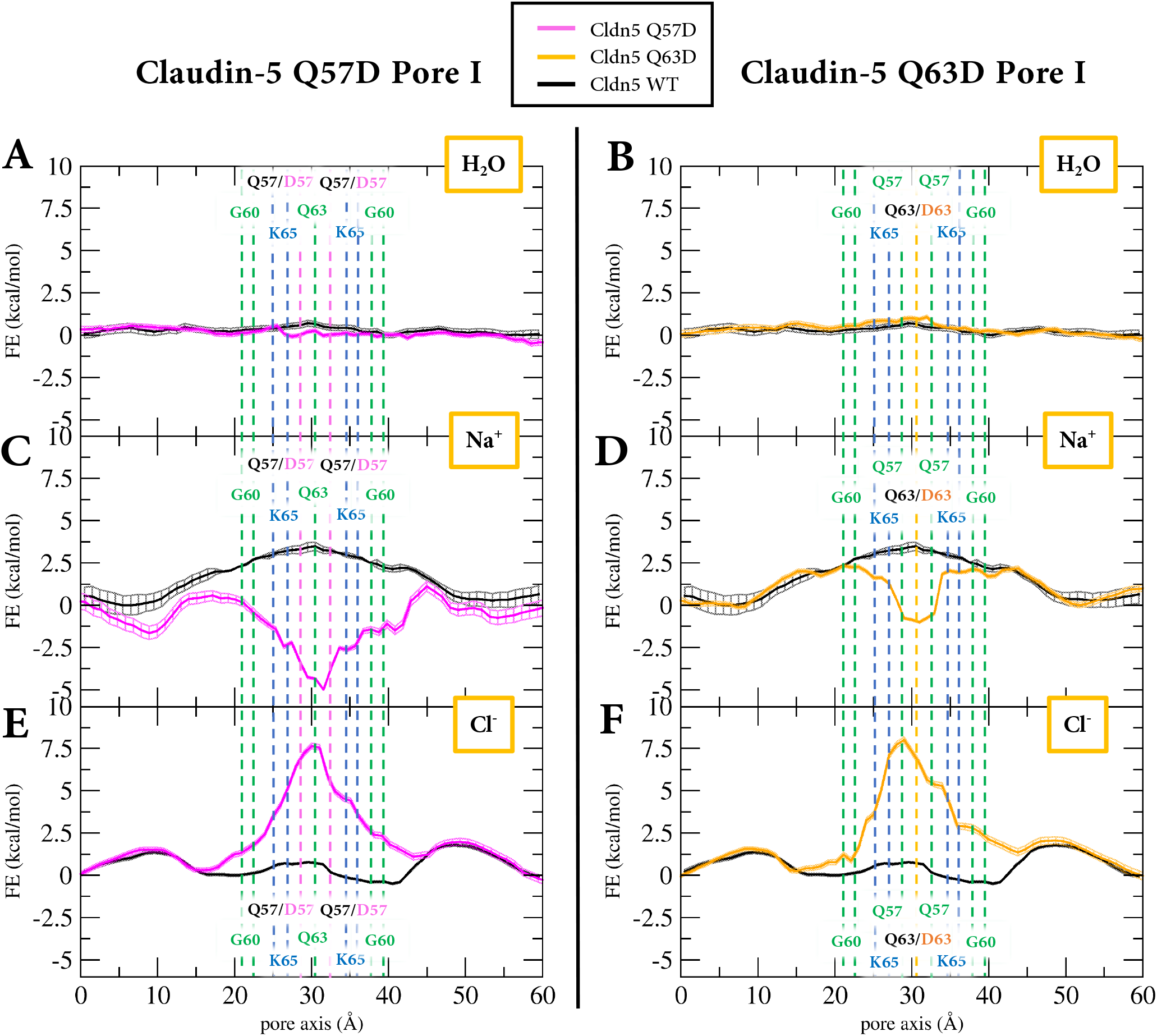
FE profiles for H_2_ O, Na^+^ and Cl^−^ translocation across the pore axis of the mutated Pore I systems. Pink and orange lines represent values for the Q57D and Q63D mutation, respectively, while black lines are for the WT systems (WT data reproduced with permission from Ref. 24, Copyright 2022 American Chemical Society). The positions of the C_*α*_ atoms of Q57(D), G60, Q63(D) and K65 are reported as dashed lines.

## Discussion

Cldn5 is an essential protein for maintaining the BBB microenvironment, thus representing a novel target in the development of molecules to efficiently permeate the barrier in a safe and reversible manner. In lack of experimental information, several structural models of Cldn5 tight junctional assemblies have been proposed, and their investigation, refinement and validation are mandatory steps to unravel the molecular basis of its function. To this goal, the recent discovery of the first Cldn5 variant associated with a neurological condition (G60R)(25) serves as an excellent bench test. The substitution, causing alternating hemiplegia in two unrelated patients, affects an amino acid between the *β*3 and *β*4 strands on ECL1, and was experimentally demonstrated to promote anion permeability in Cldn5 TJs. When inserted into two of the most studied models of Cldn5 single-pore structures, the mutation produced markedly different effects, allowing us to identify the one model displaying the experimentally observed anion-channel property. This is the so-called Pore I system, derived originally from the architecture of Cldn15 TJs proposed by Suzuki et al. (10), and first introduced for Cldn5 by Irudayanathan et al. (6).

In this model, G60 is located close to the middle of the pore cavity, and its substitution with an arginine strongly affects ion permeation. Additional simulations showed that other mutations (Q57D and Q63D, also at the center of the pore) have the effect of converting the Cldn5 complex into a cation channel. Q57 corresponds to D55 of the cation selective Cldn15, for which the validity of Pore I, and the role of the aspartic acid, have been assessed (11, 15).

The study of ionic transport in Cldn-based paracellular cavities is a quite recent and promising research field in the context of MD simulations. The results exposed in this manuscript are based on the investigation of single and isolated tetrameric pores, not considering the role of the Cldn subunits in the TJ strand that are adjacent to the pore cavity. This strategy, already employed by us (12, 13, 24) and others (6, 23), allows to considerably reduce the size of the studied systems, while retaining the key molecular determinants of TJ selectivity. Indeed, our previous work on the Cldn15 single-pore (12) revealed the same selectivity properties of a multi-pore model (15). However, the representation of the protein within a physiological environment may provide further insights by investigating the role of neighboring protomers in the strand, as various computational studies have suggested that Cldn polymerization can affect the paracellular structures (11, 14–17, 61, 62). As for the G60R mutation, in our single pore model the side-chains point towards the exterior of the cavity, while in a multi-pore structure they would penetrate the neighboring pores, possibly enhancing the electrostatic effects described here.

Another relevant element in all-atom MD simulations that could affect the results of ionic permeation events through ion channels is the choice of the force field (FF). In this study, we used the more recent version of the CHARMM FF (CHARMM36m (36–38)) FF, in agreement with all available computational works on paracellular channels (6, 7, 11– 18, 23, 24). However, recent studies (63, 64) revealed different outcomes from microsecond-long MD simulations of ion permeation through the selectivity filter of a voltage-gated potassium channel performed with CHARMM and another widely used FF, AMBER (65). Hence, future investigations of Cldn-based pores will benefit from the integrated use of multiple FFs.

In conclusion, our results provide the first *in-silico* description of the effect of a Cldn5 pathogenic mutation. They agree in full with the recently reported experimental observations (25) and allow to discern the best pore model to describe the Cldn5 assembly in the BBB. By identifying the protein-protein interface at the core of Cldn5 TJ assemblies, our computational investigation provides a step forward in the structure-based search for BBB opening agents for therapeutic treatments.

## Main abbreviations and notations

BBB: blood-brain barrier
CLDN: claudin
CV: collective variable
ECL: extracellular loop
FF: force field
FE: free energy
MD: molecular dynamics
POPC: phosphocholine
TJ: tight junction
TM: trans-membrane
US: umbrella sampling
WHAM: weighted histogram analysis method

## Author Contributions Statement

- **Conceptualization:** Giulio Alberini and Luca Maragliano.
- **Data curation:** Alessandro Berselli.
- **Formal analysis:** Alessandro Berselli, Giulio Alberini and Luca Maragliano.
- **Funding acquisition and resources:** Luca Maragliano and Fabio Benfenati.
- **Investigation:** Alessandro Berselli, Giulio Alberini and Luca Maragliano.
- **Project administration:** Giulio Alberini and Luca Maragliano.
- **Supervision:** Luca Maragliano and Fabio Benfenati.
- **Visualization:** Alessandro Berselli.
- **Writing - original draft:** Giulio Alberini.
- **Writing - review & editing:** Giulio Alberini, Alessandro Berselli, Luca Maragliano and Fabio Benfenati.

## Acknowledgments

We thank Mattia Pini and Sergio Decherchi for the kind assistance at Fondazione Istituto Italiano di Tecnologia computing center. We are grateful to Diego Moruzzo, Elisabetta Colombo and Andrea L. Benfenati for useful help. Computing time allocations were granted by the CINECA supercomputing center under the ISCRA initiative. We also gratefully acknowledge the HPC infrastructure at Fondazione Istituto Italiano di Tecnologia.

## Ethics statements

In this work, no potential identifiable human images or data are presented.

